# Validation of cell-free protein synthesis aboard the International Space Station

**DOI:** 10.1101/2023.12.06.570403

**Authors:** Selin Kocalar, Bess M. Miller, Ally Huang, Emily Gleason, Kathryn Martin, Kevin Foley, D. Scott Copeland, Michael C. Jewett, Ezequiel Alvarez Saavedra, Sebastian Kraves

**Affiliations:** Leigh High School, 5210 Leigh Ave, San Jose, CA 95124, United States; Massachusetts Institute of Technology, 77 Massachusetts Ave, Cambridge, MA 02139, United States; Division of Genetics, Brigham and Women’s Hospital, Harvard Medical School, 75 Francis St, Boston, MA 02115, United States; miniPCR bio, 1770 Massachusetts Ave, Cambridge, MA 02140, United States; Boeing Defense, Space & Security, 6398 Upper Brandon Dr, Berkeley, MO 63134, United States; Department of Chemical and Biological Engineering, Northwestern University, 2145 Sheridan Rd, Evanston, Illinois 60208, United States; Department of Bioengineering, Stanford University, Stanford, California 94305, United States

**Keywords:** cell-free protein synthesis, fluorescence, biosensor, synthetic biology, molecular biology in space

## Abstract

Cell-free protein synthesis (CFPS) is a rapidly maturing *in vitro* gene expression platform that can be used to transcribe and translate nucleic acids at the point of need, enabling on-demand synthesis of peptide-based vaccines and biotherapeutics, as well as the development of diagnostic tests for environmental contaminants and infectious agents. Unlike traditional cell-based systems, CFPS platforms do not require the maintenance of living cells and can be deployed with minimal equipment; therefore, they hold promise for applications in low-resource contexts, including spaceflight. Here we evaluate the performance of cell-free BioBits® platform aboard the International Space Station by expressing RNA-based aptamers and fluorescent proteins that can serve as biological indicators. We validate two classes of biological sensors that detect either the small molecule DFHBI or a specific RNA sequence. Upon detection of their respective analytes, both biological sensors produce fluorescent readouts that are visually confirmed using a handheld fluorescence viewer and imaged for quantitative analysis. Our findings provide insight into the kinetics of cell-free transcription and translation in a microgravity environment and reveal that both biosensors perform robustly in space. Our findings lay the groundwork for portable, low-cost applications ranging from point-of-care health monitoring to on-demand detection of environmental hazards in low-resource communities both on Earth and beyond.

**Visual graphic:** 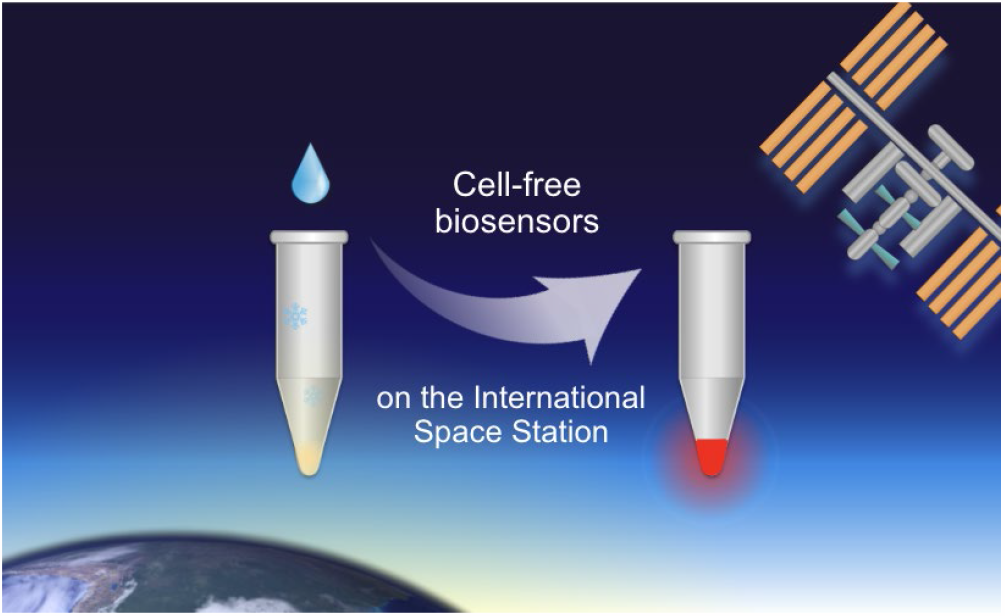

## Introduction

Advances in synthetic biology have enabled the design of versatile bioengineered technologies to address challenges in human health, biomanufacturing, environmental protection, and more (1, 2). In particular, the development of cell-free protein synthesis (CFPS) using *in vitro* gene expression platforms has allowed for these advances to take place by more efficient means outside of the traditional barrier of living systems (3-5). The applications of CFPS technology have broadened significantly in recent years, from the synthesis of protein-based vaccines and therapies at the point of care to the development of diagnostic tests for medicinally and environmentally relevant markers (6-12).

The affordable and widely applicable biotechnologies that can be developed using CFPS platforms hold immense potential in low-resource settings, including in space (13, 14). Since cell-free reactions do not contain living cells, unlike whole-cell systems, they do not need to be cultured, do not need maintenance by specialized equipment, and do not require biocontainment (15). Here, we demonstrate use of the CFPS platform BioBits® aboard the International Space Station (ISS), enabling the development of technologies that may resolve long-standing challenges in space and on Earth. BioBits® is an ideal synthetic biology tool for low-resource environments as it is not only low-cost and portable, but it is also freeze-dried for long-term stability [**Figure 1A**] (16-18).

**Figure 1:**
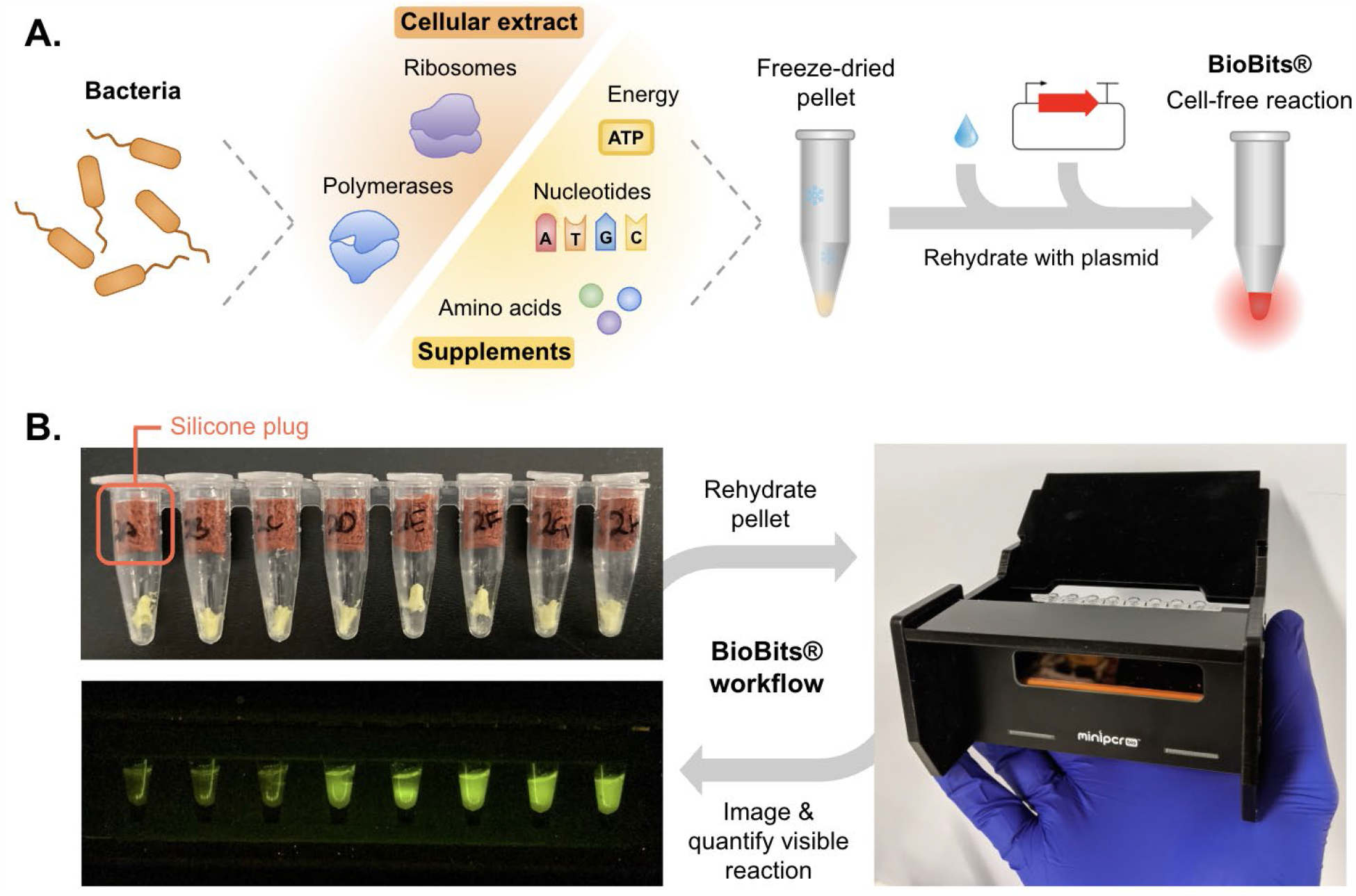
BioBits® is a tool for cell-free transcription and translation that can be coupled with the Genes in Space Fluorescence Viewer for rapid reaction monitoring. (A) BioBits® is prepared by lysing bacterial cells and lyophilizing their cellular extract along with supplements for long-term storage. When needed for use, the cell-free extract is rehydrated with the respective plasmid(s), which the cell-free machinery expresses. (B) The cell-free extract is stored in compact, portable tubes plugged with silicone stoppers to keep the lyophilized material at the bottom of the tube. This design helps ensure that the lyophilized material stays in place during micropipetting and that it doesn’t become dislodged in low gravity conditions. After rehydration with plasmids and as the cell-free reaction proceeds, fluorescent readouts can be directly visualized with the Genes in Space Fluorescence Viewer and imaged with a tablet, phone, or other capture device for quantitative analysis.

By expanding the range of lightweight synthetic biology tools aboard the ISS, we help reduce astronauts’ dependence on Earth for conducting essential research and monitoring, as there is currently a limited selection of technologies suitable for the unique demands of spacecraft relative to Earth. For example, there currently exists a strong reliance on ground-based testing facilities for monitoring pathogenic contamination aboard spacecraft (19-23). Potentially contaminated samples are periodically flown down to Earth and analyzed using complex laboratory procedures before astronauts can be informed of pathogenic contamination (19-23). Recently, on-orbit sequencing workflows coupled with bioinformatics were validated aboard the ISS, which serves as an effective method for characterizing the population distribution of bacteria (24, 25). As an alternative approach, CFPS biosensors could be developed for quick and simple detection of specific pathogenic water contaminants.

Up to now, it has been unclear how the space environment affects the performance of cell-free systems, as changes in fluid dynamics due to decreased gravity and buoyancy forces along with changes in surface tension aboard the ISS may impact enzymatic activity, substrate distribution within the liquid medium, and overall reaction mixing and convection (26-28). Here, we introduce BioBits® to the space molecular biology toolkit and evaluate its performance in microgravity. First, we qualitatively assess the two biological processes upon which CFPS applications are built, transcription and translation, by using BioBits® in its lyophilized and liquid extract forms to express RNA-based aptamers and fluorescent proteins aboard the ISS. Then, we deploy this technology towards the quantitative detection of small molecule and nucleic acid analytes using two classes of biosensors that produce fluorescent signals in response to binding to their corresponding analyte. All fluorescent signals produced through these experiments are directly monitored using the Genes in Space Fluorescence Viewer, a handheld device for real-time visualization of fluorescence, and imaged with an iPad for quantitative analysis of reaction kinetics [**Figure 1B**] (29).

In-flight validation of cell-free technology as a platform for protein expression and biosensor design may enable future developments of on-demand protein production platforms and diagnostic devices for use on long-duration space travel and in resource-limited communities. Our pipeline toward evaluating cell-free expression kinetics aboard the ISS—which uses BioBits® in complement with the Genes in Space Fluorescence Viewer—not only provides insight into how the space environment affects these biological processes, but also supports the development of versatile biotechnologies with impacts on Earth and beyond.

## Results & Discussion

### Cell-free protein synthesis

To evaluate the efficiency of cell-free systems in space, we synthesized RNA-based aptamers and fluorescent proteins using BioBits®. Although the BioBits® CFPS system is inactive when freeze-dried, the cellular machinery is able to resume transcription and translation upon rehydration. Each experiment was begun either by rehydrating the lyophilized cell-free pellet in water containing the plasmid(s) of interest or by thawing frozen cell-free extract to which plasmids had already been added.

All tubes were plugged with silicone stoppers to prevent loss of the cell-free pellet under microgravity [**Figure 1B**]. During micropipetting steps, micropipette tips were temporarily inserted between the silicone stopper and the wall of the reaction tube and then removed, allowing the silicone stoppers to remain inside the tube throughout the duration of the experiment. These experiments were repeated using liquid cell extract to determine whether freeze-drying is a suitable delivery method for BioBits® reactions to be transferred to spacecraft.

We began by rehydrating the cell-free pellet in water and Plasmid 1, which encodes the Broccoli RNA aptamer followed by the coding sequence for eforRed red fluorescent protein (RFP) [**Figure 2A**] (17, 18). In this experiment, as Plasmid 1 was transcribed, the Broccoli RNA could bind to the DFHBI fluorophore, a supplement already present in the BioBits® cell-free pellet, to form a green fluorescent complex (30). As translation of the resulting transcript occurred, RFP was synthesized, causing a gradual accumulation of red fluorescence.

**Figure 2:**
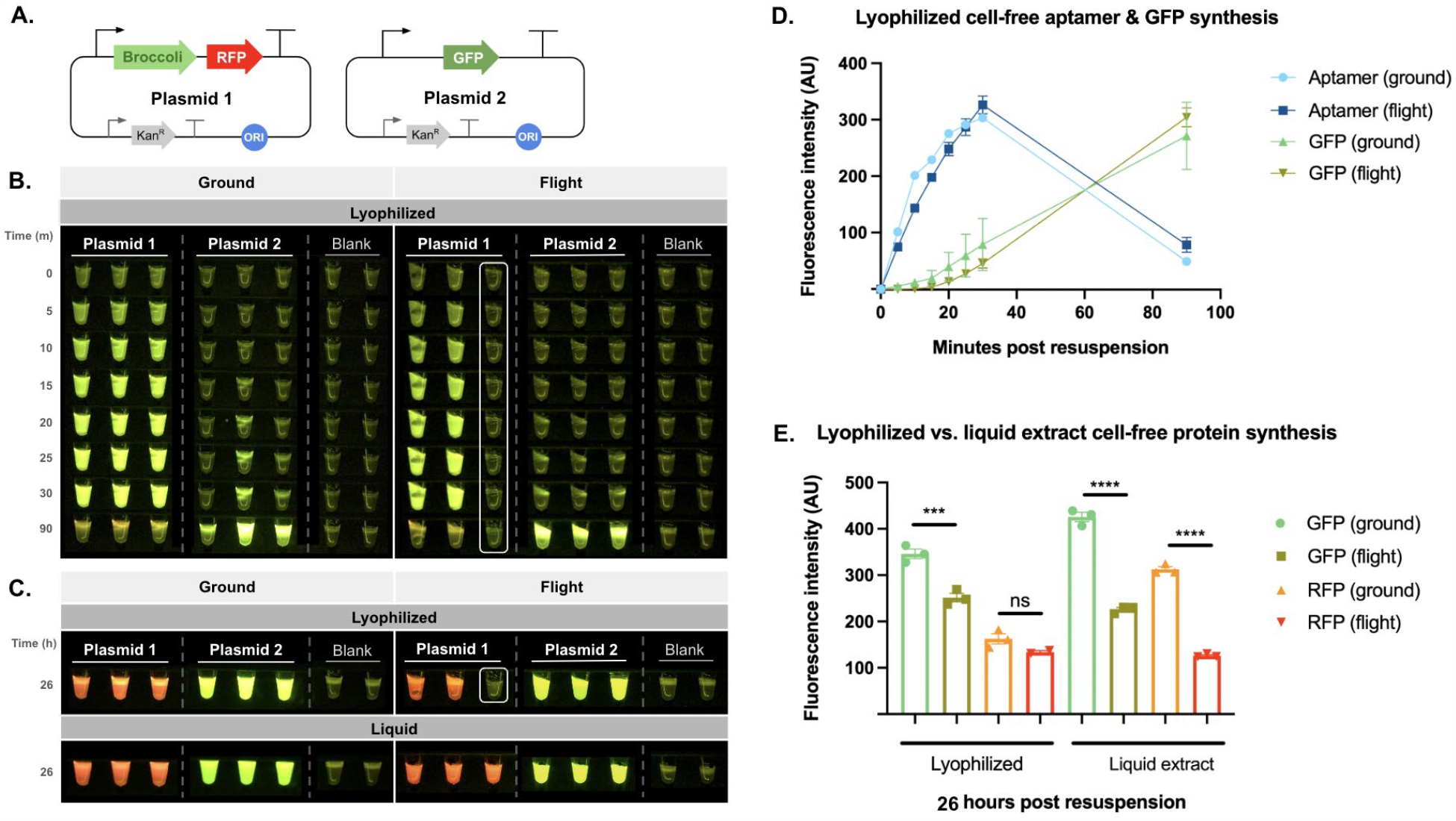
Evaluation of BioBits® transcription and translation capabilities on Earth and in space using lyophilized and liquid extract forms. (A) Two plasmid constructs were expressed by BioBits® to compare cell-free expression kinetics, Plasmid 1 encoding the Broccoli aptamer followed by RFP and Plasmid 2 encoding GFP. (B) Fluorescence images were taken at various time points following delivery of Plasmids 1 and 2 to lyophilized BioBits® reactions, showing a comparison of samples run on ground and in flight aboard the ISS. Green fluorescence is visible from aptamer transcription in reactions expressing Plasmid 1 and green fluorescence gradually emerges from GFP synthesis in reactions expressing Plasmid 2. The pellet of the sample boxed in white was not fully resuspended and thus excluded from the quantitative analysis. (C) Fluorescence images were taken after letting the cell-free reaction proceed overnight, showing RFP produced from expression of Plasmid 1 and GFP produced from expression of Plasmid 2. The pellet of the sample boxed in white was not fully resuspended and thus this sample was excluded from quantitative analyses. (D) Fluorescence intensity (arbitrary units; AU) resulting from aptamer transcription and GFP synthesis was determined through quantitative image analysis and compared between lyophilized ground and flight samples, with standard error depicted. (E) Fluorescence intensity (AU) resulting from GFP and RFP after letting the reaction proceed overnight was used to compare overall protein yield between lyophilized and liquid extract samples run on Earth and on the ISS. Data shown are mean ± SEM. Significance was calculated using 1-way ANOVA with Tukey’s multiple comparison test. ns: not-significant, ***p<0.001, ****p<0.0001

Second, to compare overall protein synthesis kinetics and yield, this workflow was repeated with Plasmid 2, which encoded free-use green fluorescent protein (GFP) variant b [**Figure 2A**] (31). As transcription and translation occurred, the synthesis of GFP caused a gradual appearance of green fluorescence. The production of the fluorescent aptamer complex (Plasmid 1) and fluorescent proteins (Plasmids 1 and 2) were visualized in the Genes in Space Fluorescence Viewer and captured using an iPad imaging system. Fluorescence intensity over time was calculated for each tube per time point and used to compare differences in transcription and translation kinetics between Earth-run and ISS-run samples, and between lyophilized and liquid extract samples [**Figure 2**].

Our findings demonstrate excellent overall performance of cell-free systems under spaceflight conditions. As observed in **Figures 2B** and **2D**, tubes with Plasmid 1 revealed similar aptamer fluorescence time courses of samples run on ground and in space. Moreover, as observed in **Figures 2C** and **2E**, Plasmid 2 readouts showed a similarly timed appearance of GFP, suggesting that translation also follows comparable time courses under both ground and flight conditions. By the 26-hour time point, substantial RFP and GFP levels were visualized in all samples, demonstrating that CFPS works robustly in both lyophilized and liquid extract forms in space. These experiments were carried out by astronauts with minimal micropipetting training, providing evidence that this method may be more accessible and easier to implement than more complex molecular workflows such as sequencing-based pipelines.

In general, fluorescence intensity tended to be higher in ground-run samples than in ISS-run samples. While this may be due to the effects of the space environment on the cell-free reaction, it may also be due to the impact of user-to-user variability on experimental results in these systems. Because the ISS is a difficult environment in which to set up highly controlled experiments, there may be inconsistencies in reaction volumes and pellet rehydration, which may cause the observed variations in resulting fluorescence intensity. In particular, one of the pellets in the lyophilized cell-free sample was not fully resuspended during rehydration with Plasmid 1 in space and was therefore excluded from analyses [**Figures 2B, C**]. Fluorescence images and further kinetic analyses of liquid extract cell-free systems over the initial 90-minute window can be found in **Supplementary Figure 1**, and indicate similar trends of expression on ground and in space. **Supplementary Figure 2** provides a comparison between liquid extract and lyophilized systems over the 90-minute window, and shows similar patterns of aptamer and GFP synthesis over time. These findings indicate that both lyophilized and liquid extract CFPS platforms are functional in space and executable by minimally trained personnel.

### Cell-free biosensor development

We next used BioBits® to test two classes of biological sensors—one that uses the Broccoli aptamer system to detect the small molecule DFHBI and another that uses a toehold switch to detect a specific RNA sequence. By testing two classes of biosensors, we sought to establish general methods of biological detection that could be repurposed toward a broader library of analytes.

To verify whether the aptamer-based system could be used as a biosensor showing concentration-dependent responses to varying amounts of its target analyte, we supplemented the lyophilized cell-free reaction with varying concentrations of the small molecule DFHBI in the presence of Plasmid 3, which encodes the Broccoli aptamer [**Figure 3A**]. Having previously established that this system was functional [**Figure 2**], we then demonstrated that it was able to detect DFHBI at varying concentrations tested in the range of 6.25 μM to 100 μM, producing a fluorescent readout positively correlated to the concentration of DFHBI [**Figure 3B, C**]. DFHBI detection was rapid and produced a visible fluorescent output within 10 minutes of beginning the reaction when compared to blank controls containing the cell-free pellet rehydrated with water only. Although flight samples tended to be brighter than ground samples, as further analyzed in **Supplementary Figure 3**, overall trends in fluorescence as a function of analyte concentration over time were preserved across both ground and flight samples as a function of time [**Figure 3C**].

**Figure 3:**
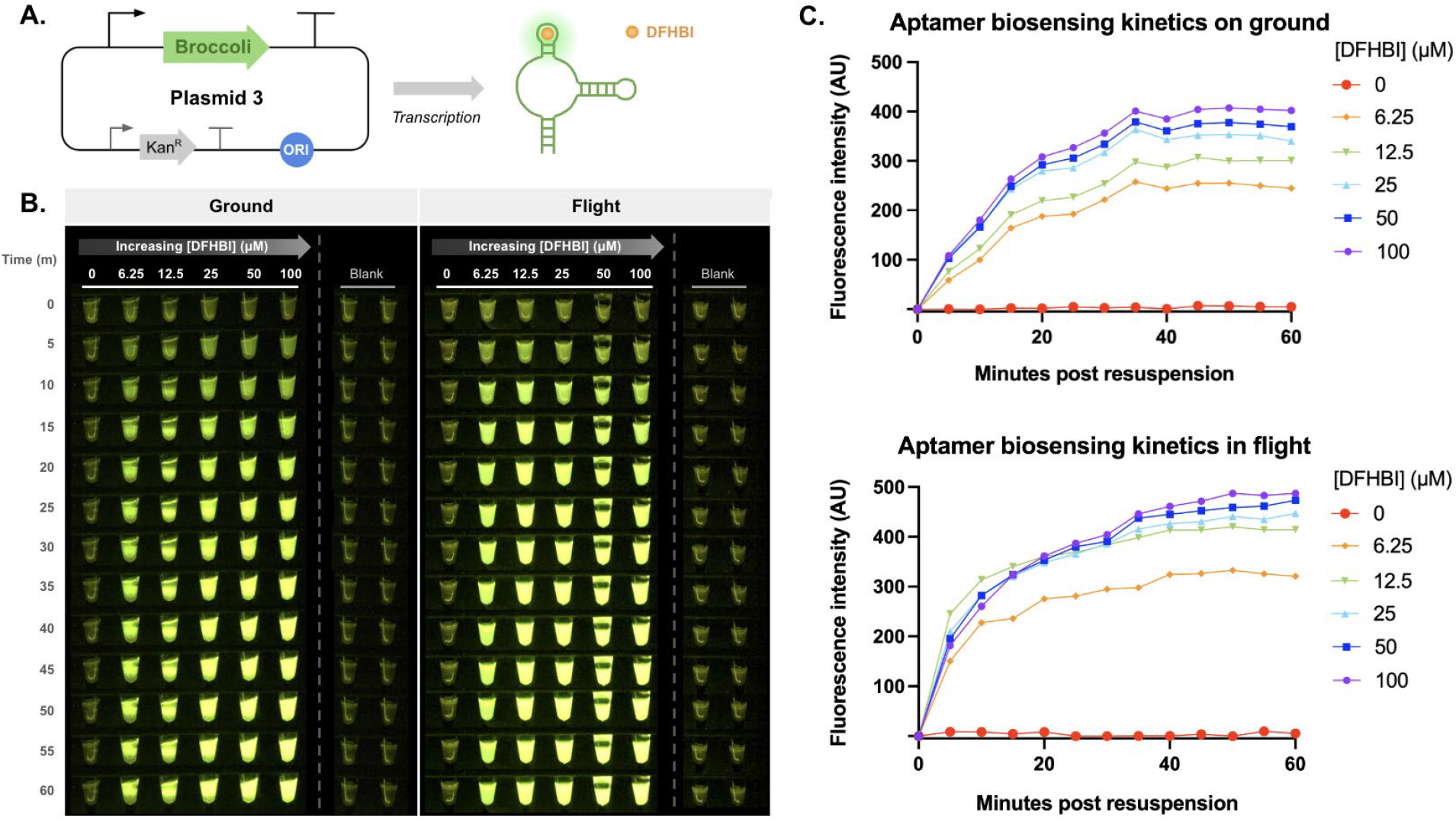
Validation of an aptamer-based BioBits® biosensor for small molecule detection on Earth and in space. (A) The Plasmid 3 construct for detecting the chemical analyte DFHBI encodes the Broccoli aptamer, which is transcribed to form an RNA that binds with DFHBI to produce green fluorescence. (B) Fluorescence images were taken at indicated time points from 0-60 minutes following rehydration of cell-free systems with Plasmid 3 and various concentrations of DFHBI, showing a comparison of samples run on ground and in flight. Green fluorescence is visible from aptamers that bind to DFHBI to produce readouts positively correlated to DFHBI concentration. (C) Fluorescence intensity (AU) from the aptamer-DFHBI complex formed in cell-free reactions was quantified at various concentrations of DFHBI for samples run on Earth and on the ISS.

We additionally validated a second class of biosensors based upon toehold switches. To operate these biosensors, we expressed Plasmid 4, a toehold switch plasmid that produces an RNA transcript which folds into a hairpin conformation, reversibly preventing ribosomal binding [**Figure 4A**] (32). This hairpin will unfold and make the ribosome binding site accessible only when a trigger RNA binds a complementary sequence contained within the hairpin. Hairpin unfolding allows for RFP translation from the downstream reporter gene only in the presence of the correct trigger RNA sequence. We evaluated this biosensor against a mismatched RNA sequence and a matching trigger RNA sequence, which were encoded in separate plasmids (Plasmid 5 and Plasmid 6, respectively) [**Supplementary Figure 4**] and co-expressed in the cell-free reaction with Plasmid 4 to determine the biosensor’s specificity [**Figure 4B**]. Plasmid 4 was also expressed with no trigger plasmid to demonstrate that no significant fluorescence signal is produced in a system without RNAs related to the hairpin sequence [**Figure 4B**]. We found that in both flight and ground samples, fluorescent readouts were only produced when the correct trigger RNA was present and that fluorescence intensity was similar between flight and ground samples [**Figure 4C**].

**Figure 4:**
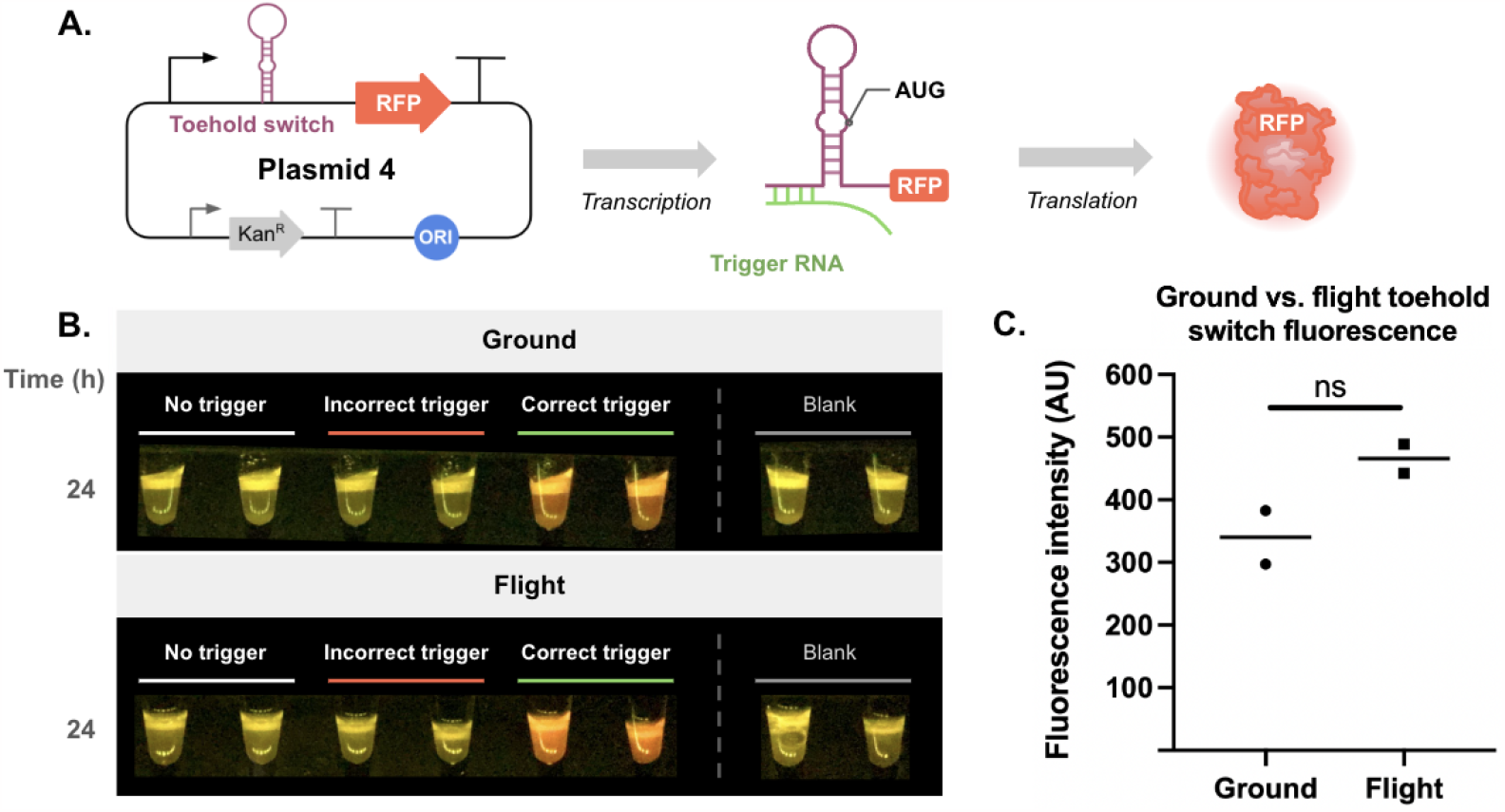
Validation of a toehold switch-based BioBits® biosensor for nucleic acid detection on Earth and in space. (A) The Plasmid 4 construct for detecting specific RNA sequences encodes a toehold switch, which is engineered to contain a complementary region to the trigger RNA sequence, followed by the reporter gene RFP. Upon transcription, the resulting RNA transcript folds into a hairpin conformation which blocks the ribosomal binding site until binding of the trigger RNA sequence, allowing for translation and synthesis of RFP. (B) Fluorescence images were taken 24 hours after rehydration of cell-free systems with Plasmid 4 expressed on its own, co-expressed with a non-specific trigger (Plasmid 5), or co-expressed with a specific trigger (Plasmid 6) sequence. In both ground and space samples, red fluorescence is only visible in the two tubes containing the correct target RNA sequence. (C) Overall fluorescence (AU) from expression of the reporter gene RFP was compared between samples run on Earth (ground) and in space.

## Conclusions

In this study, we validated that the CFPS platform BioBits® is capable of both transcription and translation in flight and successfully deployed cell-free biosensors to detect two broad classes of analytes on the ISS. Through the use of RNA aptamers and fluorescent proteins, we monitored the kinetics of cell-free transcription and translation on ground and in flight. The synthesized aptamers and proteins could all be easily detected by crew members with minimal molecular biology training using pre-existing ISS hardware, including the compact Genes in Space Fluorescence Viewer. More broadly, given the similarity of BioBits® kits to numerous crude extract-based systems, we believe these results set the stage for distributed biotechnologies based on freeze-dried, cell-free systems (33-35).

The aptamer-based biosensor produced fluorescent signals that showed a concentration-dependent response to DFHBI with a detection threshold of 10 μM DFHBI, whereas the toehold switch-based biosensor exhibited specificity for its target RNA sequence over non-target sequences and was operational at around 10 ng/μL RNA. In addition to being robustly functional on the ISS, both biosensors are low-maintenance and user-friendly; they can be stored in commercial refrigerators for months in their freeze-dried form, are simple to set up since they require minimal equipment during handling, and provide results in a rapid manner once implemented (17, 18).

Since both designs of biosensors are modular, they can be modified to detect other biochemically significant analytes, such as environmental markers, toxins, food and water contaminants, and viral RNA (36-40). To further streamline the use of these biosensors, the plasmids could be freeze-dried within the cell-free pellet, requiring just the addition of the environmental or biological sample in order to rehydrate the pellet and begin analysis. Given the extended shelf life, potential room temperature storage, and reduced weight and volume of lyophilized samples, lyophilization of the plasmid sample would be an ideal preparation mode of CFPS platforms designed for space (41-42). Additionally, arrays or multiplexed systems of lyophilized CFPS biosensors producing analyte-specific wavelengths of fluorescence could be developed for high-throughput detection of analytes. To improve the sensitivity of these diagnostic platforms, nucleic acid sequence-based amplification (NASBA), a method for rapid isothermal amplification of nucleic acid analytes, could be incorporated prior to sample analysis or plasmids could be redesigned as a cascade of plasmids containing multiple sensing modules activating a single reporter module for amplified reporter signal (5, 43).

In this study, to adapt CFPS systems for use aboard spacecraft, we plugged the cell-free reaction tubes with silicone stoppers to prevent loss of the pellet during pipetting in microgravity. While this method was effective at keeping the pellets within the reaction tube, the pellets were, on rare occasions, still able to dislodge and potentially become stuck between the silicone stopper and wall of the tube, making it challenging to fully resuspend. Thus, to further optimize BioBits® cell-free reactions for use aboard spacecraft and reduce the learning curve of pipetting around the silicone stopper, this challenge could be overcome either by adding stabilizers and additives directly into the reaction tube to allow for better adhesion of the pellet to the bottom of the reaction tube or by designing custom microtubes shaped to better hold the pellet. Alternatively, paper-based diagnostics made by freeze-drying cell-free extract on paper strips used along with dropper bottles for sample addition would reduce the need for pipetting skills and may mitigate the effect of user variability (5).

Beyond the ISS, these results may have implications in the further development of synthetic biology tools for low-resource locations on Earth, including those designed for use in remote locations and classroom settings. Cell-free diagnostics are not only inexpensive, but are also easy to transport, store, and deploy, making them well-suited for low-resource contexts, such as in viral detection platforms for places with minimal healthcare accessibility or as part of educational platforms for classroom demonstrations (16-18, 44-48). Hence, as this synthetic biology tool continues to advance, cell-free platforms may pave the way for biological solutions to societal problems—both on and off planet Earth.

## Methods

### General template design and preparation

pET28c F30-2xdBroccoli18 was a gift from Samie Jaffrey (Addgene plasmid #66843). pUS252b (containing fuGFPb) was a gift from Nicholas Coleman (Addgene plasmid #191831). pJL1-eforRed, pNP1 eforRed trigger, pNP1 eforRed toehold switch, and pNP1 tdTomato trigger were a gift from James Collins (Addgene plasmids #106320, #107357, #107353, and #107354 respectively). pET28c F30-2xdBroccoli18 and pJL1-eforRed were used to make Plasmid 1 by cloning the F30-2xdBroccoli aptamer into pJL1-eforRed upstream of the eforRed gene via Gibson assembly. fuGFPb was used to make Plasmid 2, pET28c F30-2xdBroccoli18 was used to make Plasmid 3, and the eforRed toehold switch was used to make Plasmid 4. tdTomato trigger and eforRed trigger were each cloned into separate pJL1 backbone plasmids to create the mismatched trigger (Plasmid 5) and matching trigger (Plasmid 6), respectively, for the eforRed toehold switch. Cloning and plasmid propagation were performed using NEB Turbo (New England Biolabs; C2984H) competent *E. coli* cells.

### In-house crude cell-free extract preparation and lyophilization protocol

Cell extract was prepared as described previously (49). Briefly, *E. coli* BL21 Star (DE3) cells (Thermo Fisher Scientific) were grown in 2× YTPG media at 37°C at 250 rpm. T7 polymerase expression was induced at an OD600 (optical density at 600 nm) of 0.6 to 0.8 with 1 mM isopropyl-β-d-1-thiogalactopyranoside. Cells were harvested in mid-exponential growth phase [OD600 = ∼2 to 3], and cell pellets were washed three times with ice-cold Buffer A containing 10 mM tris-acetate (pH 8.2), 14 mM magnesium acetate, 60 mM potassium glutamate, and 2 mM dithiothreitol, flash-frozen, and stored at −80°C. Then, cell pellets were thawed and resuspended in 1 mL of Buffer A per 1 g of wet cells and sonicated in an ice water bath. Total sonication energy to lyse cells was determined by using the sonication energy equation for BL21-Star (DE3) cells, [Energy] = [[volume (μl)] − 33.6] * 1.8 − 1. A Q125 Sonicator (Qsonica) with 1/4” diameter probe at a frequency of 20 kHz was used for sonication. An amplitude of 50% in 10 second on/off intervals was applied until the required input energy was met. Lysate was then centrifuged at 12,000 relative centrifugal force (rcf) for 10 min at 4°C. The supernatant was flash-frozen and stored at −80°C until use.

The cell-free reaction mixture consisted of the following components: 1.2 mM adenosine 5′-triphosphate; 0.85 mM each of guanosine-5′-triphosphate, uridine 5′-triphosphate, and cytidine 5′-triphosphate; L-5-formyl-5,6,7,8-tetrahydrofolic acid (34.0 μg ml−1; folinic acid); E. coli transfer RNA mixture (170.0 μg ml−1); 130 mM potassium glutamate; 10 mM ammonium glutamate; 12 mM magnesium glutamate; 2 mM each of 20 amino acids; 0.33 mM nicotinamide adenine dinucleotide; 0.27 mM CoA; 1.5 mM spermidine; 1 mM putrescine; 4 mM sodium oxalate; 33 mM phosphoenolpyruvate; 10 μM DFHBI 1T (Tocris; 5610), which was included in all samples except for the aptamer-based biosensor samples; and 27% (v/v) of cell extract (11, 12, 35).

Prepared cell-free reactions were either aliquoted into 8-strip 0.2 mL microtubes at volumes of 20 μL and flash-frozen in liquid nitrogen for use in liquid extract cell-free experiments or aliquoted into 8-strip 0.2 mL microtubes at volumes of 25 μL, flash-frozen in liquid nitrogen, and lyophilized overnight for use in lyophilized cell-free experiments. All Earth-run samples were stored in a -80°C freezer until use. All ISS-run samples were launched to the ISS at -80°C and stored in the Minus Eighty Laboratory Freezer for ISS (MELFI) until use.

### Reaction packaging for spaceflight use

To prevent the lyophilized reactions from leaving the reaction tubes, particularly in microgravity conditions, silicone plugs were inserted into the reaction tubes before capping the tubes. Circular plugs were cut from 0.25” thick low compression set silicone sponge (Stockwell R10480S) using a ¼” diameter biopsy punch. Each tube was plugged with one silicone plug.

### Reaction reconstitution and characterization

The reactions were thawed at room temperature before use. To reconstitute the cell-free reactions, the user slightly compressed the silicone plug to the side to allow the micropipette tip through and dispense the reagents into the tube. When the micropipette tip was removed from the tube, the silicone plug decompressed back to its original state to keep the reaction in the tube. The reactions were reconstituted with nuclease-free water containing the relevant plasmid(s) and incubated at room temperature in the Genes in Space Fluorescence Viewer. Per each strip of reaction tubes, two tubes served as blank controls and contained the already hydrated cell-free pellet (in the case of liquid extract cell-free experiments) or the rehydrated cell-free pellet (in the case of lyophilized cell-free experiments).

The final concentration of plasmid in the cell-free reaction was 25 ng/μL of Plasmid 1 (encoding the Broccoli aptamer followed by RFP), 5 ng/μL of Plasmid 2 (encoding GFP), 17 ng/μL of Plasmid 3 (encoding the aptamer-based biosensor), or 12 ng/μL of Plasmid 4 (encoding the toehold switch-based biosensor). For the biosensor experiments with Plasmid 4, the cell-free pellet was also supplemented with either 12.6 ng/μL of Plasmid 5 (encoding the incorrect trigger RNA) or 12.6 ng/μL of Plasmid 6 (encoding the correct trigger RNA).

### Imaging settings and analysis

Images of the reactions were taken at the specified time points using the Yamera application on an iPad Pro, which was connected to the Genes in Space Fluorescence Viewer by its Viewer Adapter. Imaging settings for comparisons of lyophilized and liquid extract cell-free transcription and translation were [Aspect Ratio 4:3; Focus locked at 0.1; Shutter at 1/20; ISO at 1000±10 during the first 90 minutes of imaging and changed to 500±10 for next-day imaging; Tint at -150; and Temp at 4179±10], imaging settings for the aptamer-based biosensor were [Aspect Ratio 4:3; Focus locked at 0.1; Shutter at 1/20; ISO at 1000±10; Tint at -150; and Temp at 4179±10], and imaging settings for the toehold switch-based biosensor were [Aspect Ratio 4:3; Focus locked at 0.1; Shutter at 1/20; ISO at 1760±10; Tint at -100; and Temp at 8000±10].

All images were analyzed using ImageJ, through which images were converted to 8-bit binary form and measurements of integrated density and mean fluorescence were taken of each tube (50). For each strip of 8 tubes, these measurements were taken of a constant, set area for ease in normalization to the 2 blank tubes on each strip (containing the already hydrated or rehydrated cell-free pellet). The corrected fluorescence was then calculated as follows: integrated density – (0.5 * area * (mean fluorescence of 1st blank + mean fluorescence of 2nd blank)). Finally, initial fluorescence values were subtracted from all other fluorescence readings for background subtraction.

## Acknowledgments

We thank Jesse Rusk from miniPCR bio for devising the silicone stopper configurations, Melissa Boyer, Linda Gibson, and Agha M. Abbas from The Boeing Company for flight integration support, and NASA Astronaut Robert Hines for completing this experiment aboard the ISS. This work was supported in part by New England Biolabs and the ISS National Laboratory. M.C.J. gratefully acknowledges the National Science Foundation (CBET: 1936789), the Army Research Office (W911NF-22-2-0246; W911NF-22-2-0210), and the Department of Energy (DE-SC0023278).

