## Supplementary Info for "Validation of cell-free protein synthesis aboard the International Space Station"

### Supporting Information for
### ***Validation of Cell-Free Protein Synthesis Aboard the International Space Station***

Selin Kocalar, Bess M. Miller, Ally Huang, Emily Gleason, Kathryn Martin, Kevin Foley, D. Scott Copeland, Michael C. Jewett, Ezequiel Alvarez Saavedra, Sebastian Kraves*

*Corresponding author contact:


- Liquid extract cell-free protein synthesis …
  1. Liquid extract aptamer and GFP synthesis kinetics SI-2
  2. Liquid extract vs. lyophilized aptamer and GFP synthesis SI-3
- Cell-free biosensor development …
  3. Ground vs. flight aptamer-based biosensing SI-4

#### Liquid extract cell-free protein synthesis

##### 1) Liquid extract aptamer and GFP synthesis kinetics

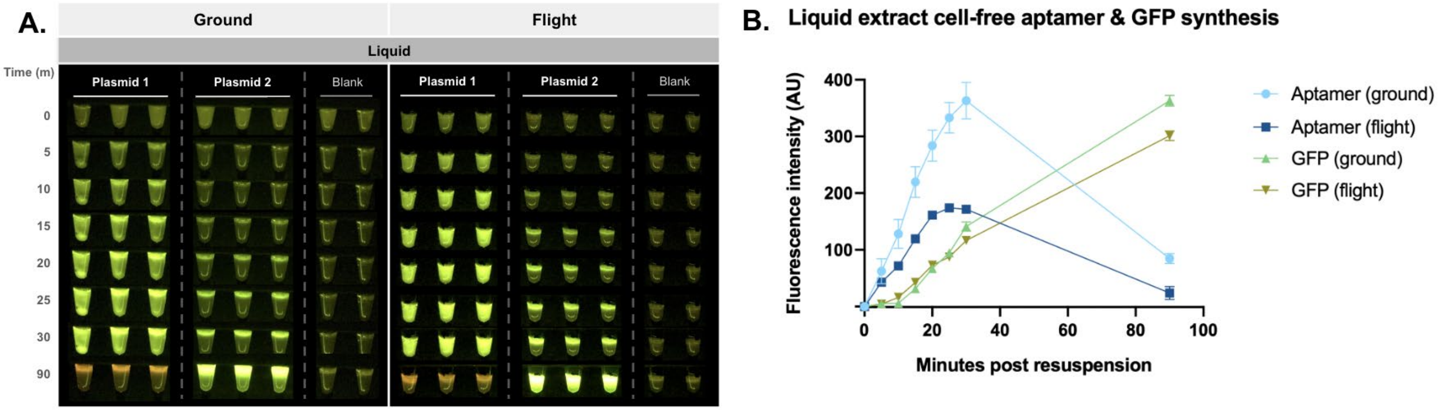


**Figure S1: Evaluation of liquid extract BioBits® transcription and translation capabilities on Earth and in space.** (A) Fluorescence images were taken at indicated time points following delivery of Plasmids 1 and 2 to liquid extract BioBits® reactions, showing a comparison of samples run on ground and in flight aboard the ISS. Green fluorescence is visible from aptamer transcription in reactions expressing Plasmid 1 and green fluorescence gradually emerges from GFP synthesis in reactions expressing Plasmid 2. (B) Fluorescence intensity (AU) resulting from aptamer transcription and GFP synthesis was determined through analysis of fluorescence images and compared between liquid extract ground and flight samples, with standard error depicted. Data shown are mean ± SEM.

#### 2) Liquid extract vs. lyophilized aptamer and GFP synthesis

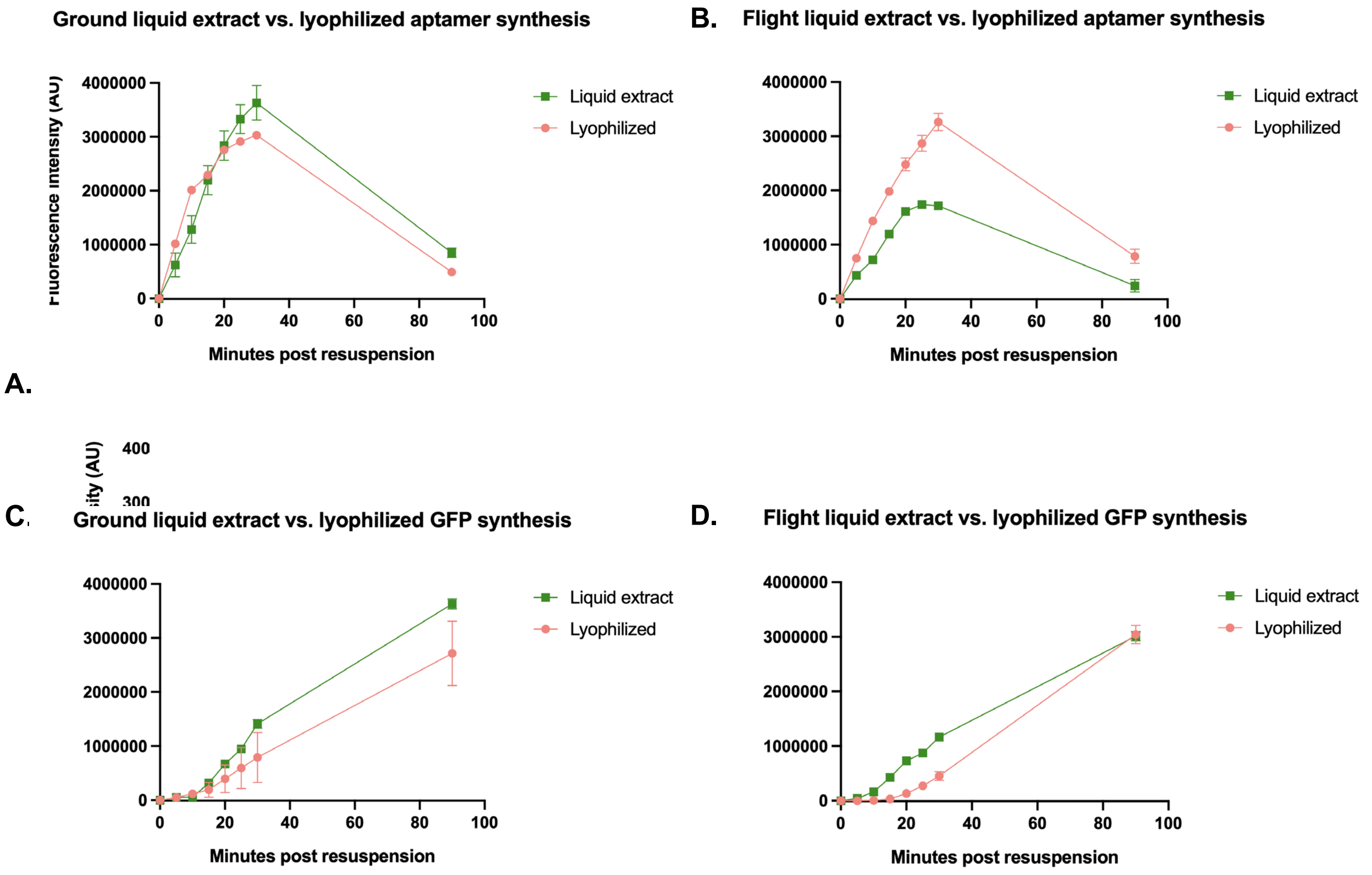


**Figure S2: Ground and flight comparisons of liquid extract vs. lyophilized aptamer and GFP synthesis, with standard error depicted.** (A) Fluorescence intensity (AU) resulting from GFP synthesis (Plasmid 2) was compared between liquid extract and lyophilized ground samples. (B) Fluorescence intensity (AU) resulting from GFP synthesis was compared between liquid extract and lyophilized flight samples. (C) Fluorescence intensity (AU) resulting from aptamer transcription was compared between liquid extract and lyophilized ground samples. (D) Fluorescence intensity (AU) resulting from aptamer transcription was compared between liquid extract and lyophilized flight samples. Data shown are mean ± SEM.

#### Cell-free biosensor development

##### 3) Ground vs. flight aptamer-based biosensing

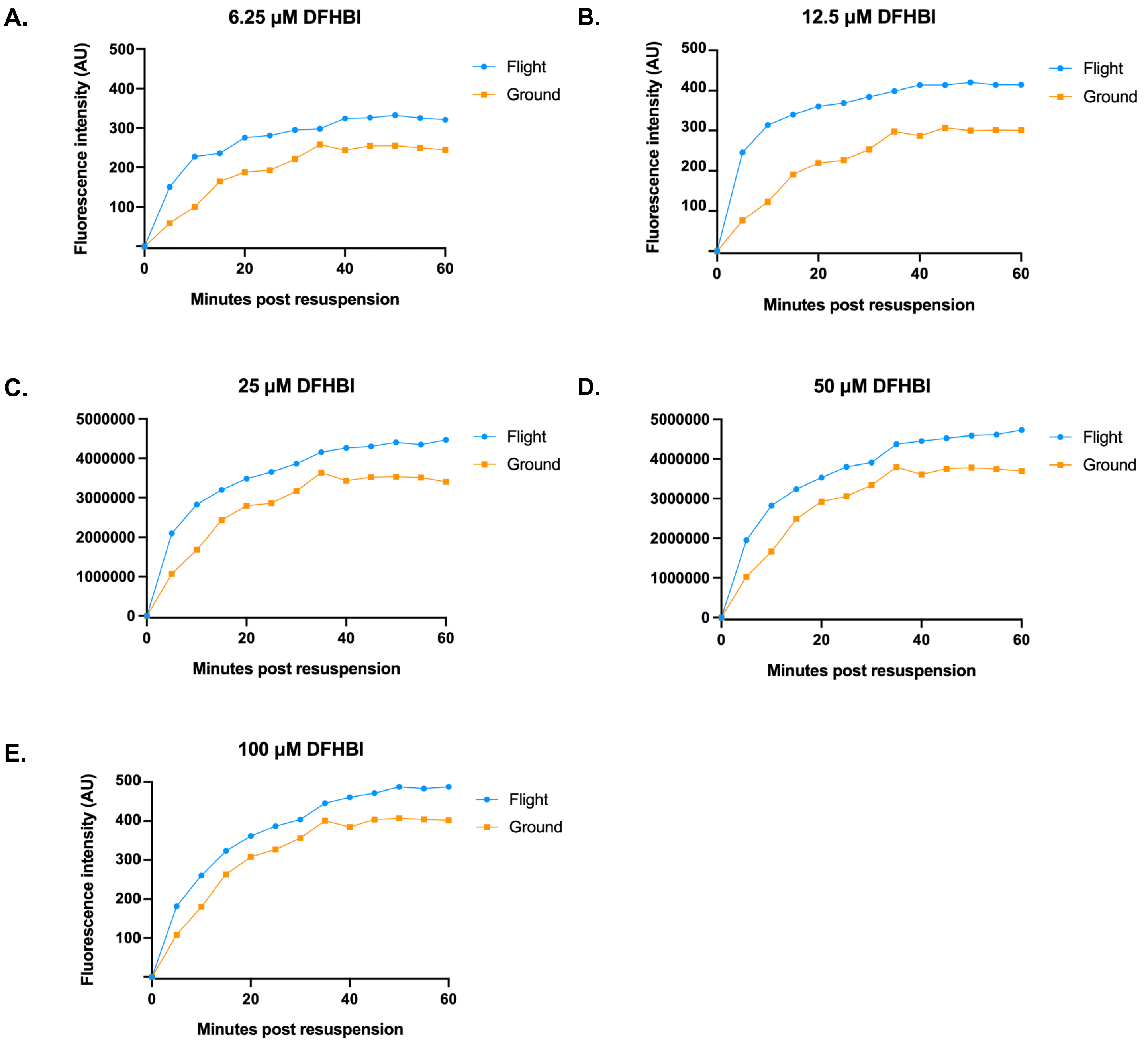


**Figure S3: Ground and flight comparisons of aptamer-based cell-free biosensing at various concentrations of the analyte DFHBI.** Fluorescence intensity (AU) resulting from the aptamer-DFHBI complex formed in cell-free reactions was compared between samples on Earth and in space at DFHBI concentrations of (A) 6.25 µM, (B) 12.5 µM, (C) 25 µM, (D), 50 µM, and (E) 100 µM.

###### 4) Ground vs. flight aptamer-based biosensing

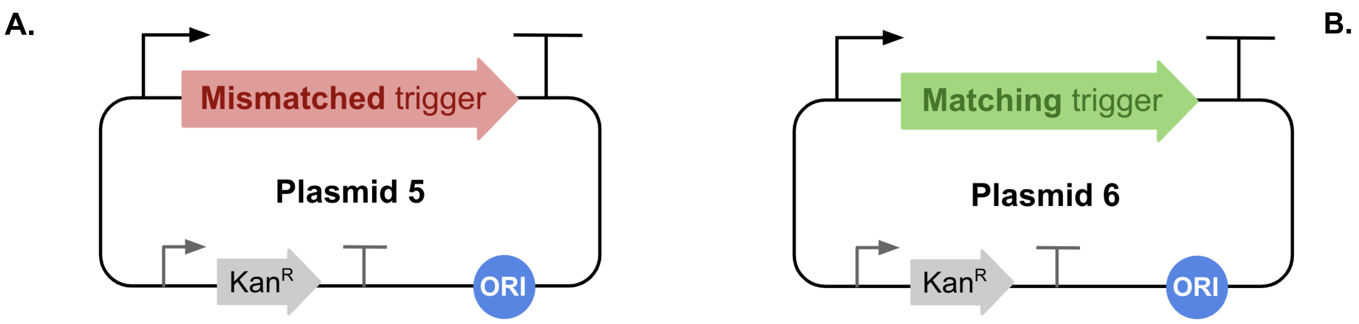


**Figure S4: RNA trigger plasmid constructs for the toehold switch-based biosensor.** (A) Plasmid 5, which encodes the mismatched RNA trigger. (B) Plasmid 6, which encodes the matching RNA trigger.
